# Clinical and Cost Effectiveness of Two “New” Lynch Syndrome Case Finding Protocols in Endometrial Cancer Population Contrasted with the IHC-based Protocol

**DOI:** 10.1101/611541

**Authors:** James M. Gudgeon, Michael W. Varner, Mia Hashibe, Marc S. Williams

## Abstract

**Purpose:** To investigate the effectiveness and costs of two Lynch syndrome screening protocols among endometrial cancer (EC) patients and compare to an immunohistochemistry (IHC)–based protocol.

**Methods:** Analytic models were developed to represent the two protocols: a brief cancer family history questionnaire (bFHQ) and direct-to-sequencing of the germline. Data from reviews of published literature, augmented by local data and expert opinion were used to populate the model representing the number of women diagnosed in the U.S. in 2018. Multiple analyses employing simulation modeling were performed to estimate a variety of clinical and economic outcomes.

**Results:** Under conditions considered here to be plausible, the bFHQ is expected to miss 58% (min./max. = 24 to 80%) of LS index cases, a direct-to-sequencing protocol to miss 30% (min./max. = 6% to 53%), and the IHC-based protocol based on previous analysis 58% (min./max. = 33 to 80%). When direct costs of testing and genetic counseling are added to the models, the total screening program costs for the bFHQ protocol are substantially lower at all sequencing price points than the other protocols. For example, at the low end of the sequencing price point (ie, $250), the total cost of screening programs for IHC, bFHQ, and sequencing are $22.9 million, $5.3 million, and $13.8 million, respectively. The best estimate of the break-even price of sequencing, when the cost of the program is equivalent between the IHC and sequencing protocols, is about $635.

**Conclusion:** The bFHQ and direct-to-sequencing LS screening protocols are more effective and cost effective at identifying LS index cases than the standard IHC-based protocol under the conditions represented in our models. These estimates of various outcome metrics of the three LS index case finding protocols may help stakeholders make decisions of the use of limited healthcare dollars.

## Introduction

In our first simulation model (submission to *Gynecologic Oncology* in progress), we made the case that Lynch syndrome (LS) screening programs starting with the immunohistochemistry (IHC) test will miss between 33 to 80 percent of the LS index cases present in a population of unselected endometrial cancer (EC) cases. We also identified, through sensitivity analyses, the relative importance of model variables on the primary outcome. The biggest single failure factor is the high rate of women who do not give consent to be sequenced after a positive screen (and passed the rule-out test, methylation of the *MLH1* promoter) thus do not get tested. Following the consent rate, the sensitivity and specificity the screening test itself, the IHC test, are the next most influential factors on index case finding, but orders of magnitude smaller than consenting. And there are many other steps in this protocol where the protocol can fail which, collectively, results in the high rate of failure to identify index cases (best estimate of about 58%).

In the past few years there has been a switch in sequencing technologies from Sanger methodologies to what is often called “next generation sequencing” (NGS).^62^ NGS is based on high-throughput, massively parallel (computerized-) processing technologies and different methods for detecting nucleotides (the building blocks of DNA), that has led to a substantial reduction in the price of DNA sequencing.^63^ The cost for a commercially available sequencing test for familial cancer syndrome genes, including the LS genes, is now approaching that of the IHC test. Sequencing, by either of these two methodologies, is the diagnostic and reference test for LS. This is the final test in the IHC-based LS screening protocol but has also been proposed as the one-and-only test for identifying the presence of index cases in patients with LS-associated cancers, primarily endometrial and colorectal cancer, and in a subpopulation of ovarian cancer. Up until recently (2017), a direct-to-sequencing protocol for identifying LS index cases has been considered too expensive, based on several cost effectiveness analyses and reviews,^15,16,64^ at least in colorectal cancer populations. No valid cost effectiveness analyses have made this determination in EC populations. However, the recent, precipitous drops in the cost of sequencing has brought increased attention to this approach. For this reason, we will explore a direct-to-sequencing protocol in this study, including all the steps in such a protocol where we believe there is meaningful potential for failure, as in our previous modeling (submission in progress).

The second screening test of interest is a simple, 4-item questionnaire of personal and family cancer history developed in Canada (Eiriksson et al 2015).^19^ This brief family history questionnaire (bFHQ) was developed and validated on a population of unselected EC patients treated at a tertiary hospital in Toronto, Ontario. In their population, this simple screen identified 100% (N=87, 95% confidence interval: 59–100%) of the Lynch syndrome index cases identified by sequencing, with reasonable specificity (about 76%). It has yet to be validated in other populations. However, it is promising, especially given the potential for cost savings with a simple questionnaire. The major advantages of bFHQ screening are that it is easy to perform and essentially free (depending on how collection and scoring are administered and “costed”).

The answers to the bFHQ questions could be obtained and scored by a clinician during routine care for these patients, office staff, or recorded by the patient themselves, as was done in the Canadian study. The primary potential disadvantage of this approach is that the patients may not know the answers to the questions at the time of asking or may answer them incorrectly; though the results from the Canadian study (ie, 100% sensitivity for LS index cases) suggests this is not problematic. Moreover, in the 2-3 years since this study was conducted multiple commercial software platforms have become available whereby patients can either fill out (cancer) family history via a web portal prior to the clinic visit or do this in the clinic with the help of a digital device. Appropriately trained clinicians can follow-up to improve the veracity of the patient-provided answers to the questions. Regardless of who and how the four questions are answered, the performance of the bFHQ requires validation in the setting in which it is being considered.

The deficiency in published cost effectiveness studies on LS screening in EC populations is the failure to compare screening protocols to the option of “no screening”. Thus, we don’t yet know whether LS screening in this population is “cost effective” in an absolute sense.

The lack of absolute cost effectiveness data for Lynch syndrome screening in EC populations, whether targeted to subsets of EC populations (eg, <70 yoa) or unselected, has not prevented medical specialty societies and others from recommending Lynch syndrome screening in EC populations. A joint recommendation by the American College of Obstetrics and Gynecology/Society of Gynecologic Oncology (2014) encourages tumor testing by IHC (or microsatellite instability, MSI) on all EC cancers in women <60 years old.^65^ More recently, several authors have called for screening in all (unselected) women with EC.^8,47^ LS screening based on age cut-offs has consistently demonstrated failure to find a substantial number of LS index cases,^55,66,67^ and there is no reason to think this would be different in EC populations.

Based on the above, the aim of this study was to examine the relative efficiency of two protocols that recent evidence and simple logic would suggest offer the potential for both superior LS index case finding and cost effectiveness, compared to the current standard for LS screening, a tumor/IHC-based protocol. We will assess affordability by calculating total screening program costs for each screening protocol and the break-even cost of sequencing.

## Methods

For this study, we developed two decision analytic models of a virtual cohort of women diagnosed with EC who were screened for LS using the two (re-) emerging protocols of interest; ie, the Canadian questionnaire (bFHQ) and direct-to-sequencing. The simulated cohort was composed of women diagnosed with EC who could be screened for LS using an IHC-based protocol. The cohort size is based on the number of new diagnoses of uterine cancer expected in the U.S. in 2018 (approximately 63,000), multiplied by 0.92 (92%), the percent of uterine cancer that are “endometrial” [16]. This results in a cohort of 58,000 new cases of EC annually. No other exclusion criteria were applied, including age at diagnosis.

The two models included all the steps we determined to have potential for failure (see Figure 1 and Table 1). The structure of the model representing the bFHQ was constructed in a manner consistent with the Eiriksson study and how we would expect it to be implemented.^19^ Similarly, the structure of the direct-to-sequencing-only protocol was constructed based on discussions with local clinicians. Both of these protocols have fewer steps and tests than an IHC protocol--a function of fewer tests, simpler requirements for those tests, and fewer hand-offs (see Figure 1). The perspective of the analysis was of healthcare systems. Models were constructed in Excel^©^ software (Microsoft, Inc., Redmond, WA) and sensitivity analyses were performed with @Risk software (Palisade, Inc., Ithaca, NY), an Excel “add-on”. Models in these formats are available by request.

**Table 1.** Model Variables and Parameters for the bFHQ Protocol

Model Variables and Parameters for the bFHQ Protocol
| Item | Parameter | Base-Case | Range | Source |
| --- | --- | --- | --- | --- |
| 1 | Prevalence of LS in population | 5.0% | fixed | targeted review of literature |
| 2 | EC patients who agree to and successfully complete bFHQ | 90% | 75-95% | based on Eriksson study<sup>19</sup> and local expert opinion |
| 3 | Sensitivity of bFHQ screen | 90% | 75-95% | based on Eriksson study<sup>19</sup> |
| 4 | Specificity of bFHQ screen | 76% | 72-80% | based on Eriksson study<sup>19</sup> |
| 5 | Consent for sequencing | 85% | 75-95% | local expert opinion |
| 6 | Sequencing ordered, blood drawn and sent | 97.5% | 95-99% | local expert opinion |
| 7 | Sequencing successful & reportable | 97.5% | 95-99% | estimate based on limited literature |
| 8 | Sensitivity of Sequencing test | 99.6% | 99-99.9% | expert opinion, targeted review of literature<sup>15,30,35</sup> |
| 9 | Specificity of Sequencing test | 99.6% | 99.0-99.9% | targeted review of literature<sup>15,30,35</sup> |
| 10 | cost of bFHQ screen | \$25 | fixed | best estimate |
| 11 | Cost of sequencing | variable | \$250-1,000 | author knowledge of commercial market |

**Figure 3.1.**
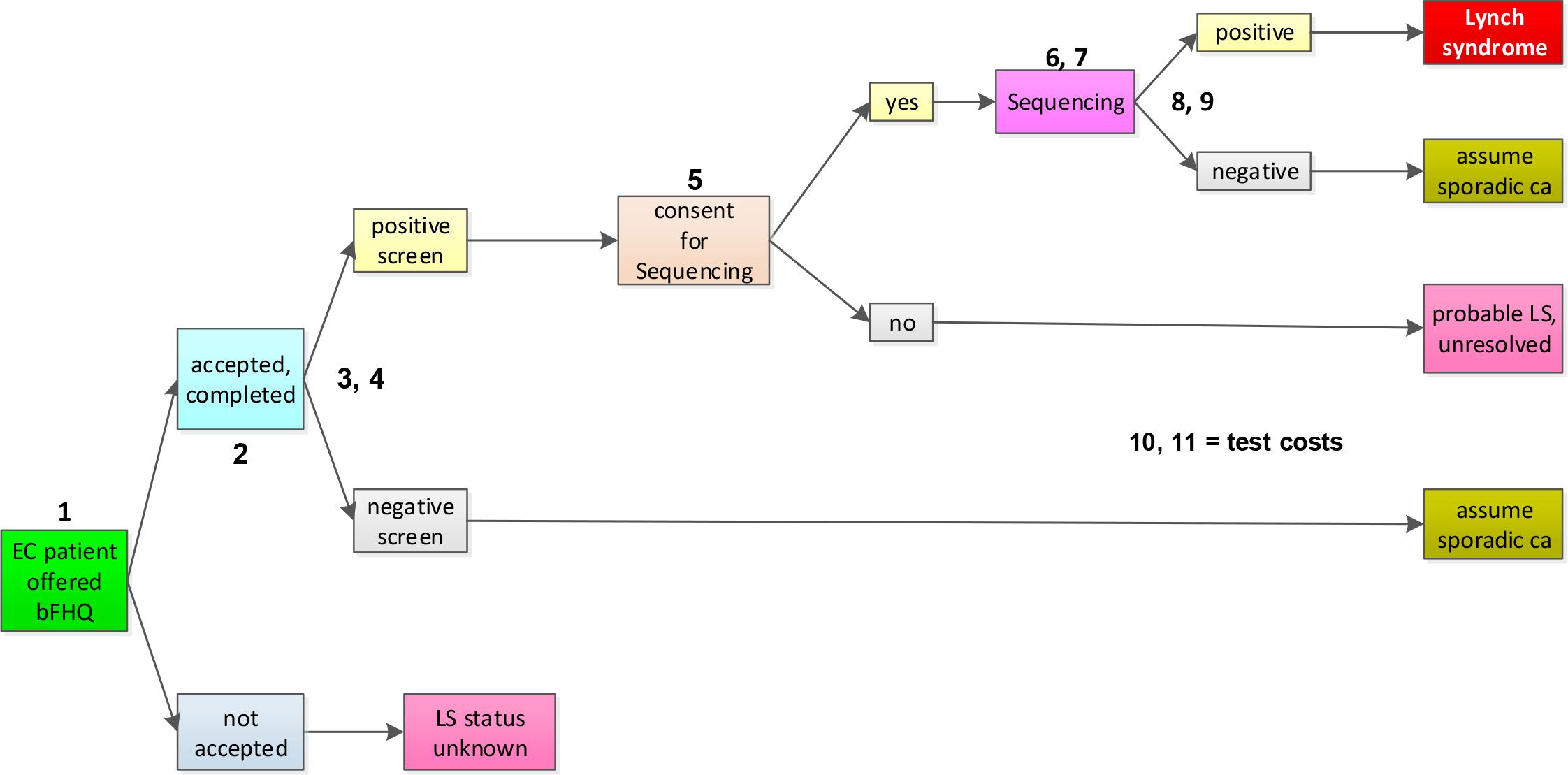
Diagram of the bFHQ protocol.

Test performance characteristics of the bFHQ were taken from the Eiriksson study, and sequencing characteristics were assumed to be the same as in the IHC protocol previously constructed (and these variables were, in fact, linked between the models). We used the 95% confidence interval of the sensitivity of the bFHQ as reported by Eiriksson and colleagues for the plausible range of this variable. To account for the negative correlation known to exist between the sensitivity and specificity values of tests,^68,69^ but for which we could identify no data to support estimations of a specific correlation values, we assigned a value of −0.75 for this model parameter (based on our previous modeling efforts, in press). Model variables and the values used to populate them are provided in Tables 1 and 2.

**Table 2.** Model Variables and Parameters for the Sequencing Protocol

Model Variables and Parameters for the Sequencing Protocol
| Item | Parameter | Base-Case | Range | Source |
| --- | --- | --- | --- | --- |
| 1 | Prevalence of LS in population | 5.0% | fixed | targeted review of literature |
| 2 | Consent for sequencing | 90% | 75-99% | local expert opinion |
| 3 | Sequencing ordered, blood drawn and sent | 98% | 97-99% | local expert opinion |
| 4 | Sequencing successful & reportable | 98% | 95-99% | conservative estimate based on limited literature |
| 5 | Sensitivity of Sequencing test | 99.5% | 99-99.9% | targeted review of literature<sup>15,30,35</sup> |
| 6 | Specificity of Sequencing test | 99.5% | 99.0-99.9% | targeted review of literature<sup>15,30,35</sup> |
| 7 | Cost of sequencing | variable | \$250, 500, 1000 | Author knowledge of commercial market |

The current study added several new outcomes to those reported from our previous model of the IHC-based protocol, enabled by the addition of costs to the screening protocols. The primary clinical outcome is percentage of LS index cases identified and missed by each protocol. Secondary clinical outcomes include number of LS cases expected to be missed and percent of LS cases identified expected to be false positives.

There two primary economic outcomes are total costs to conduct a LS screening program for a cohort of EC patients the size of the number of women diagnosed with EC in 2018, and the cost-per-LS index case-detected in the virtual EC cohort, a measure of testing efficiency. The secondary econometric metric is break-even cost of sequencing when the total program cost of the IHC-based protocol is approximately equivalent to total program costs of a direct-to-sequencing protocol.

Sensitivity analyses were performed to explore the effect that the plausible ranges of each variable in the models had on model outcomes. This included multiple one-way sensitivity analyses to identify the variables with the most influence in the two new protocols.

We conducted these analyses applying Monte Carlo simulations using @Risk software (Palisade, Inc., Ithaca, NY) as an add-on to Excel software. Secondarily, we conducted ‘what if’ analyses to explore the impact on economic outcomes as the cost of sequencing was changed from a high of $1000, approximating the high end of pricing looking forward, to a low of $250, where the current lowest market price is hovering.

## Results

The results from our models are shown in Tables 3 and 4 and represent the expected outcomes in a virtual cohort representing all women diagnosed with EC in the U.S. in 2018.

**Table 3.**
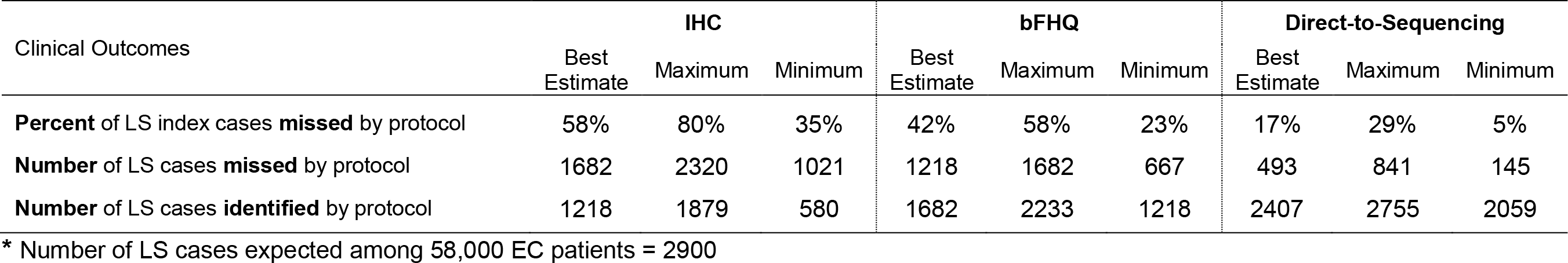
Range of Estimates of Key Clinical Outcomes

| Clinical Outcomes | IHC |  |  | bFHQ |  |  | Direct-to-Sequencing |  |  |
| --- | --- | --- | --- | --- | --- | --- | --- | --- | --- |
|  | Best Estimate | Maximum | Minimum | Best Estimate | Maximum | Minimum | Best Estimate | Maximum | Minimum |
| **Percent** of LS index cases **missed** by protocol | 58% | 80% | 35% | 42% | 58% | 23% | 17% | 29% | 5% |
| **Number** of LS cases **missed** by protocol | 1682 | 2320 | 1021 | 1218 | 1682 | 667 | 493 | 841 | 145 |
| **Number** of LS cases **identified** by protocol | 1218 | 1879 | 580 | 1682 | 2233 | 1218 | 2407 | 2755 | 2059 |

\* Number of LS cases expected among 58,000 EC patients = 2900

**Table 4.** Key Economic Outcomes of All Three Lynch Syndrome Screening/Testing Protocols at Base-Case

| Economic Metrics | Total Program Cost and Cost Effectiveness to Screen the EC Cohort When Sequencing is Priced at: |  |  |  |  |  |  |  |  |
| --- | --- | --- | --- | --- | --- | --- | --- | --- | --- |
| | **\$1,000** | | | **\$500** | | | **\$250** | | |
|  | IHC | bFHQ | Seq. | IHC | bFHQ | Seq. | IHC | bFHQ | Seq. |
| Total dollars of testing + GC time for screening program (in millions \$) | 24.402 | 13.267 | 50.679 | 22.989 | 7.480 | 25.258 | 22.258 | 4.586 | 13.080 |
| Dollars per LS index case detected (\$) | 20,245 | 6,982 | 20,262 | 19,034 | 3,936 | 10,232 | 18,428 | 2,414 | 5,244 |

Multiple one-way sensitivity analyses across the two models yielded the tornado graphs of Figure 2 and 3. As illustrated in Figure 2, there is no dominant factor for the bFHQ; rather there are four factors that drive most of the variability in outcome, given the assumptions made in our model. Figure 3 illustrates that the rate of consent for sequencing, with its wide plausible range, is the dominant threat to LS case finding in the direct-to-sequencing protocol, as is the case with the IHC-based protocol (from previous model).

**Figure 3.2.**
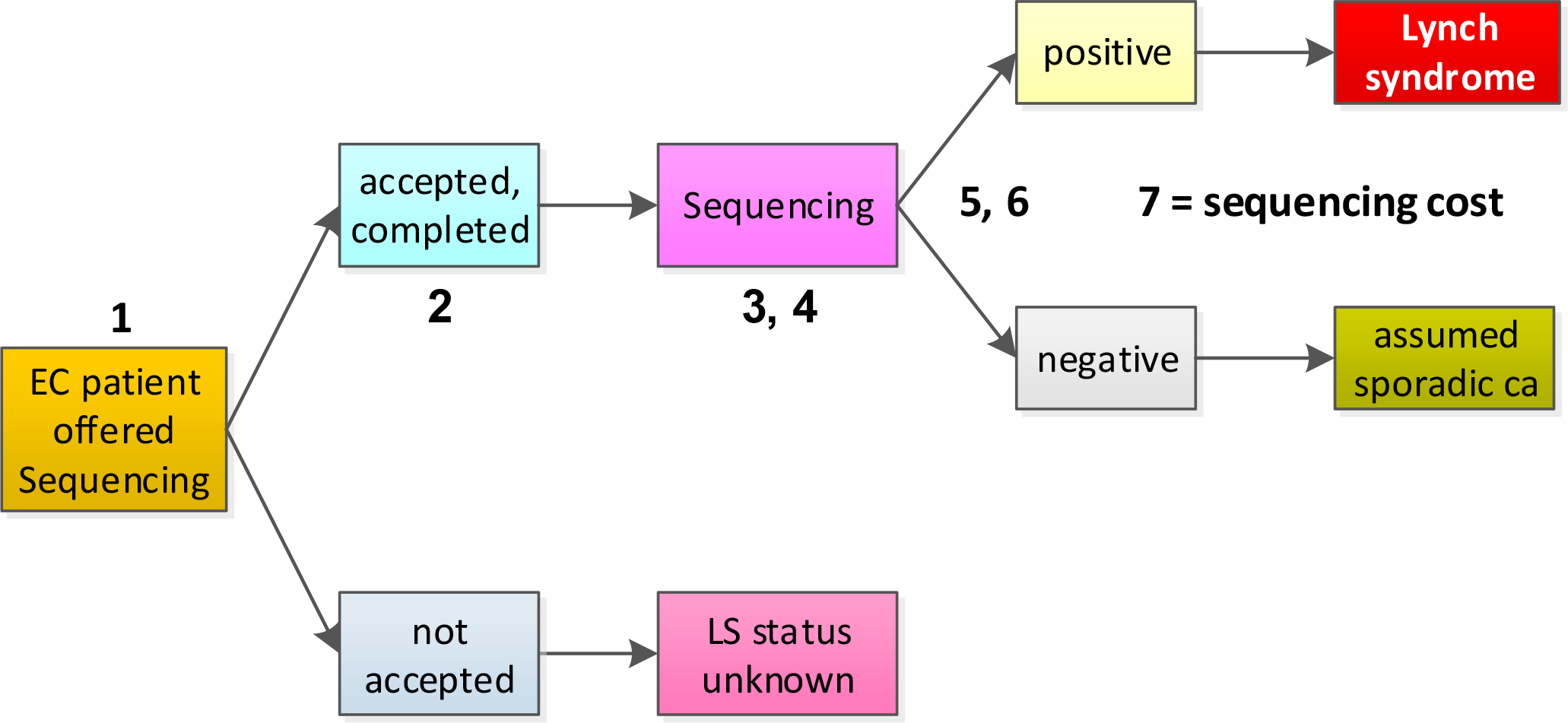
Diagram of the direct-to-sequencing protocol.

**Figure 3.3.**
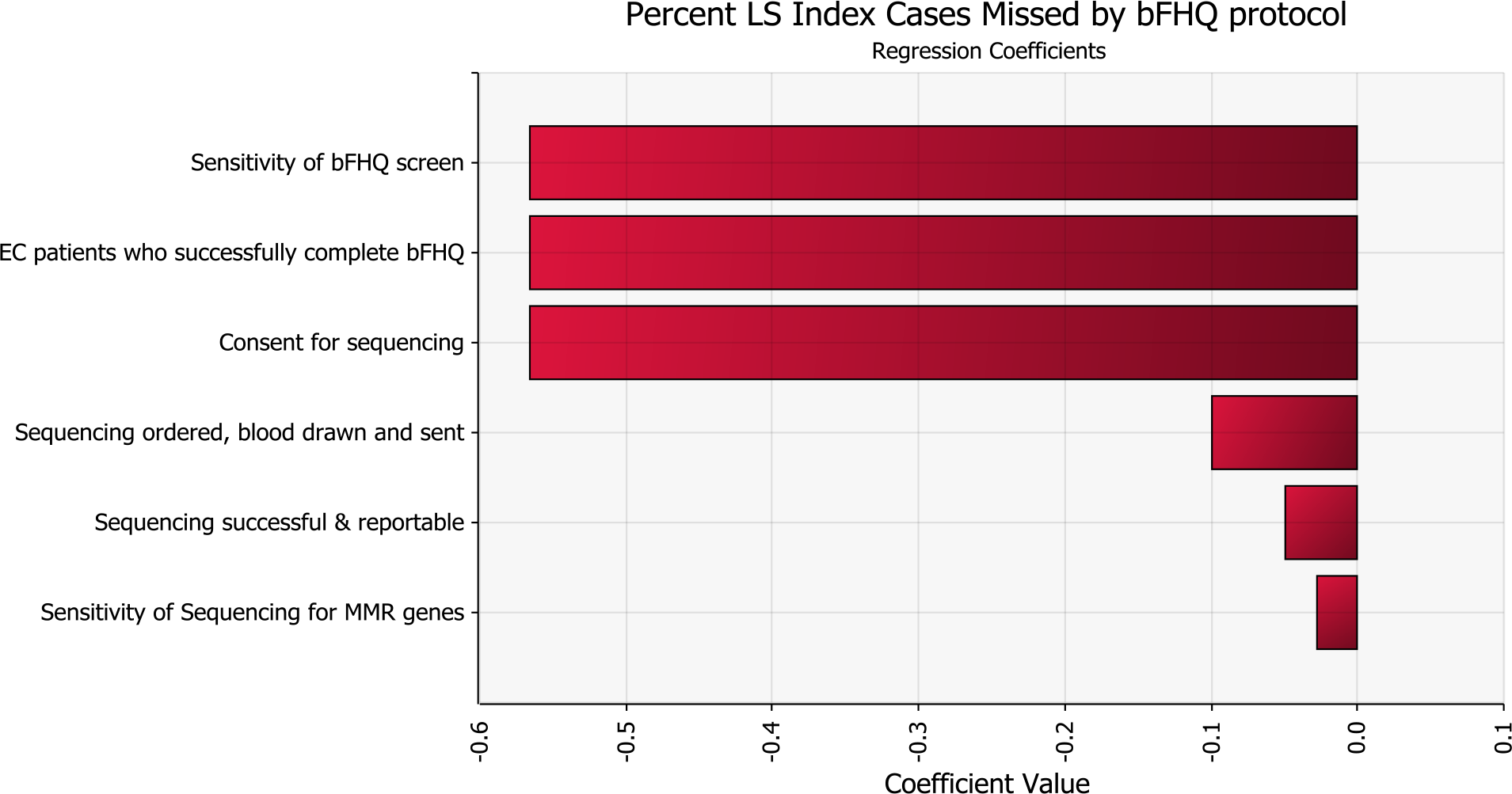
Tornado graph of the relative influence of variables on LS index cases missed by the bFHQ protocol.

**Figure 3.4.**
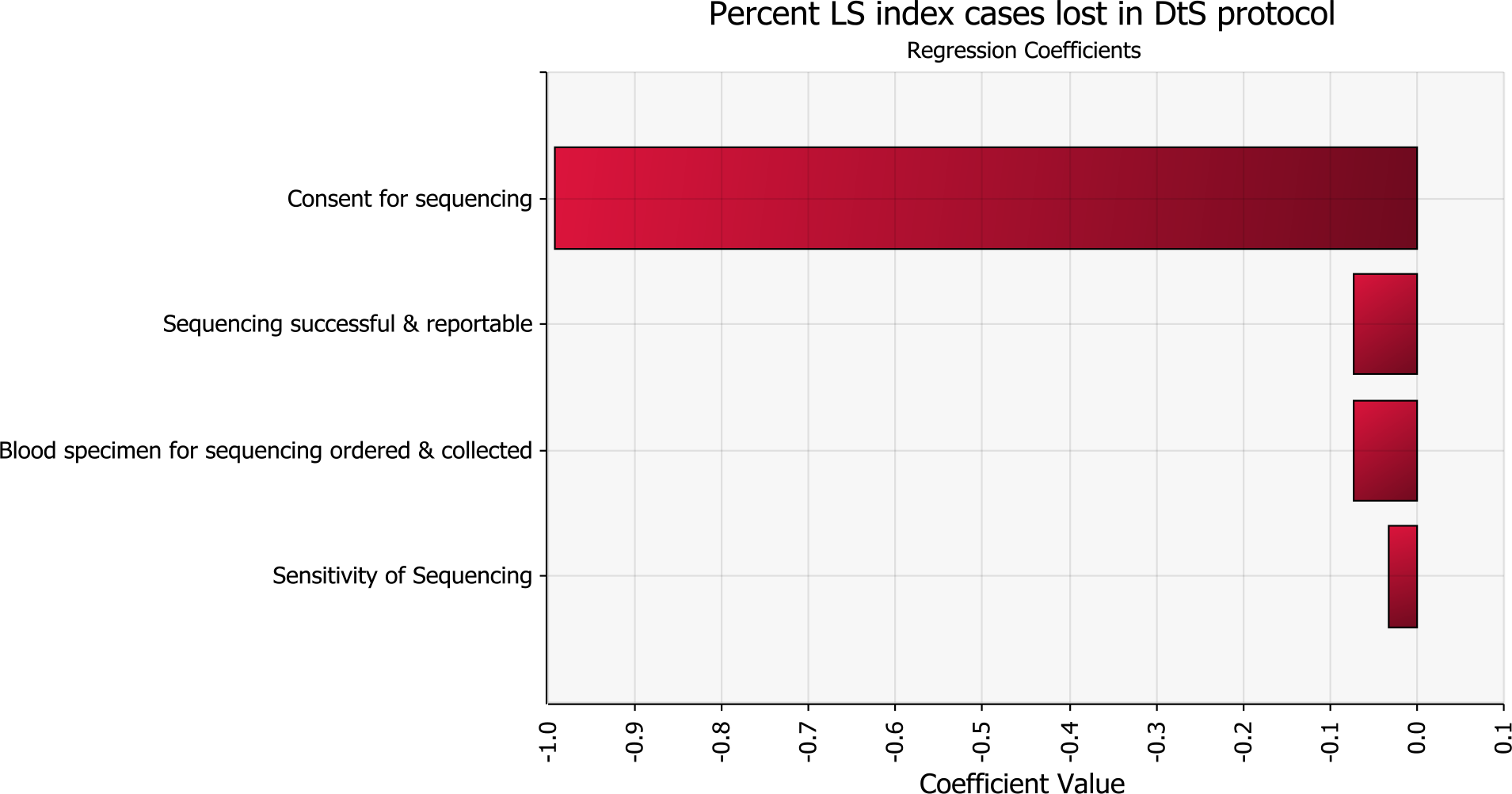
Tornado graph of the relative influence of variables on LS index cases missed by direct-to-sequencing protocol.

Table 3 compares the best estimates and minimum and maximum estimates (range) of several clinical outcomes of interest, across all three screening/testing protocols. These best estimates for failure to identify LS index cases are 42% (min./max. = 23-58%) for bFHQ, 17% (min./max. = 5-29%) for direct-to-sequencing, and 58% (min./max. = 35-80%) for IHC. The table also provides the expected numbers of missed and identified cases, out of the 2900 presumed to be present in the cohort.

Table 4 provides estimates of the total direct costs of a LS screening/testing program and the cost-per-LS index case-detected in the virtual EC cohort, when the cost of sequencing was valued at $1000, $500, and $250 across the three protocols.

The best estimate of break-even cost of sequencing when the total program costs of the IHC-based protocol is equivalent to total program costs of a direct-to-sequencing protocol is about $445 (we excluded the bFHQ from this analysis).

## Discussion

We developed two decision models to investigate the plausible ranges of failure rates in LS index case finding and various other outcomes, including economic, in the bFHQ and direct-to-sequencing protocols. We compared the outcomes from these models with those previously generated on the IHC-based screening protocol for LS cases (publication in submission) to which we added costs to the model.

The one-way sensitivity analyses explored the relative impact of different factors of each protocol. For the bFHQ, there is no dominant factor; rather there are four influential ones: consent for sequencing, sensitivity of the bFHQ, specificity of the bFHQ, and percent of bFHQ’s that are interpretable. We added the interpretability variable late in the study as we discovered this to be a challenge in a local trial (Intermountain) even when scored by licensed genetic counselors who specialize in cancer genetics.

With the direct-to-sequencing protocol, outcome variability is all about the consent rate, even more so than the IHC protocol (from previous model). This is due to the narrow plausible ranges of the other (non-consent) variables in contrast to the wide range of consent entered into the model. The latter reflects the lack of published data on this issue and variability encountered between enrolling gyn-oncology physicians in our local study. One of the central issues underlying consent in this context is whether a high or even 100% consent rate is desired--from an ethical and legal viewpoint. That is, do patient and medical provider communities agree about the ideal outcome in this context; e.g. maximal LS index case finding versus fully informed patient consent and choice making. We will tackle this issue in a subsequent paper.

The primary model outcome of percent of LS index cases missed can be interpreted as expected index case finding under “average” conditions of test protocols, derived from variables populated with best estimates and statistical averages. That is, if a mix of healthcare systems were to apply the three LS case finding protocols to all 58,000 of the women in the U.S. diagnosed with EC in 2018, we expect our “best estimates” to reflect the expected outcomes. The minimum and maximum outputs correspond to most optimistic and most pessimistic testing conditions in the same virtual population. We also point out that the direct-to-sequencing protocol, under the most optimistic, plausible testing conditions is expected to identify 95% of index cases (5% missed). This compares to 77% identified (23% missed) for the bFHQ and 65% identified (35% missed) for the IHC protocol.

The total LS case index finding program costs provide estimates of the costs of the three different protocols at each price point of sequencing ($1000, $500, $250) in our fictional cohort of average screening/testing programs. These estimates illustrate that bFHQ and direct-to-sequencing protocols are heavily influenced by changes in sequencing costs whereas the change in program costs for the IHC protocol is small. That is, the drop in prices due to the benefits from next generation sequencing will have big impacts on the bFHQ and direct-to-sequencing but little on the current standard protocol, IHC. This, in itself, may be another reason to consider switching from the IHC protocol to direct-to-sequencing or, IF it can be validated in local settings, the bFHQ. This begs the question of what sequencing price point will yield equivalent costs for a direct-to-sequencing versus IHC-based case finding protocol; that is, the break-even point. In our models that break-even price point is about $445. To add another dimension to this issue of value, we provide estimates of the cost per index case identified, which illustrates the huge increases in case finding *efficiency* in a direct-to-sequencing protocol compared to IHC.

All these estimates, clinical and economic, can and should be contextualized by examining variable inputs, as shown in Tables 1 and 2. The challenge of such as exercise reinforces the necessity of a thorough understanding of the structure of a given simulation model as well as the numbers assigned to model variables to help decide whether and how it is applicable to a local setting. Much of the uncertainty built into our models can be reduced if not eliminated by customizing the models with local data, thereby providing more accurate ‘best estimates’ and narrow min./max. ranges (or CIs). For this reason, we encourage interested stakeholders to request a digital copy of our models and populate the variables with site-specific numbers, or create a model reflecting local conditions.

We remind readers that these differences in rates of failure and success index case finding cascade through families as the multiple family members who are LS carriers do not get the chance to undergo testing for the family LS mutation, with subsequent follow-up preventive or treatment interventions.

It is clear that the IHC protocol cannot compete with a direct-to-sequencing protocol in terms of clinical effectiveness. However, its cost effectiveness has previously been determined to be unacceptably poor,^15–17,71^ albeit at yesterday’s sequencing costs (eg, $2,000-3,000). At today’s and tomorrow’s sequencing cost, the superior sensitivity of the direct-to-sequencing protocol offers no only its superior index case finding, but also offers substantially better cost effectiveness.

This study has several limitations. The first limitation is the lack of empiric data to provide robust estimates of the consenting phase of both the bFHQ and direct-to-sequencing protocols, influential variables in all protocols (on clinical outcomes). The consenting process is likely to be substantively different from consenting as it is currently practiced in the IHC-based protocol which, in the U.S., is nearly always managed and administered by genetic counselors, with a “non-directive” approach.^40,72^ Consenting in the bFHQ and direct-to-sequencing protocols, as envisioned in this study, will be managed and administrated by gynecologic oncologists in the setting of EC, who may be more “directive”.^73,74^ We addressed the uncertainty of consenting rates in this setting by applying relatively wide ranges around these variables, but higher values/ranges than consenting in the IHC protocol (which has a strong literature base justifying its relatively low consent rates).

The second limitation is our representation of LS screening as universal; that is, offered to all women with EC, as opposed to targeted population. We realize that other institutions take somewhat different approaches to LS screening (eg, screen only EC patient <70 years old, use of the MSI screen). Any such differences in approach would influence, perhaps substantially, the performance of these protocols in those settings and generate different estimates of the metrics reported here.

The third possible limitation is our use of a higher LS prevalence rate (5%) than what some investigators have identified in their studies. We based this estimate on recent studies that obtained complete ascertainment of LS index cases in their populations, in contrast to most older studies that only performed the gold standard diagnostic test on cases that screened positive. If the actual rate is substantially different then the absolute values for study outcomes will vary; however, the relative values will vary little if at all.

Finally, some readers may be quick to reject the validity of the bFHQ since it has not been validated outside of the Canadian population where it was developed, and various studies have verified the challenges associated with reliable family history taking in similar clinical settings.^75^ However, the potential of this screening approach is too great to not include it in modeling studies. We addressed these valid concerns by accounting for the uncertainty in performance of the screening protocol in the modeling. We are testing this protocol internally at Intermountain.

In the direct-to-sequencing and IHC protocols (previously modeled), consenting is the dominant variable in the identification of index cases. In the bFHQ there are four protocol factors that share dominant influence on outcome, one of which is consenting. As we discussed in our previous publication (in press, Gudgeon et al, *Gynecologic Oncology*), there are many reasons for poor consent rates in this context and little to no reason to think they will improve, at least not in the paradigm of the current IHC-based protocol.

A break-even price point for sequencing of $445 seems well within near-term reach, thus changing from an IHC-based to direct-to-sequencing case finding protocol seems inevitable if not imminent. The huge difference in testing efficiency as measured by cost per LS case detected further demonstrates the potential benefit of this change. Again, we remind readers that these numbers will vary substantially from program to program thus justifying efforts to estimate them under local conditions. Moreover, the bFHQ protocol holds great promise but does require validation in any given population/institution.

Based on our analyses, the time seems to have arrived for stakeholders responsible for LS patient populations and their families, to take a hard look at how well current screening programs are performing, and dollars spent. We hope our analyses, and perhaps use of our Excel-based models to fit local conditions, can help others sort out important questions about implementation or continued application of LS case finding.

After deciding which screening protocol is the best fit for a particular institution, all institutions must overcome known hurdles to complete the entire care pathway. These hurdles are cascade testing in families of index cases, and provision of evidence-based interventions that reduce morbidity and mortality from LS-associated cancers. These activities require exceptional coordination of services, further complicated by the complexities of insurance coverage. Effective implementation of these activities following effective LS case finding are necessary to reach the goal of substantially reducing the morbidity and mortality associated with LS-associated cancers.

